# Notch signaling regulates *Akap12* expression and primary cilia length during renal tubule morphogenesis

**DOI:** 10.1101/760181

**Authors:** Malini Mukherjee, Ishara Ratnayake, Madhusudhana Janga, Eric Fogarty, Shania Scheidt, Justin Grassmeyer, Jennifer deRiso, Indra Chandrasekar, Phil Ahrenkiel, Raphael Kopan, Kameswaran Surendran

## Abstract

Alagille syndrome patients present with loss of function mutations in either *JAG1* or *NOTCH2*. About 40-50% of patients have kidney abnormalities, with multi-cystic, dysplastic kidneys being one of the more frequent kidney defects. Additionally, gain-of-function mutations in *NOTCH2* are associated with cystic kidneys in Hajdu-Cheney syndrome patients. Conditional inactivation of *Notch1, Notch2*, or *RBPJ* within the nephrogenic lineage impairs nephrogenesis and produces proximal tubule cysts in mice. How perturbations in Notch signaling cause renal tubular cysts remains unclear. Here we have determined that inhibition of Notch signaling in the kidney increases *Akap12* expression. Ectopic expression of Akap12 in renal epithelia results in abnormally long primary cilia similar to those observed in Notch-signal-deficiency. Both loss of Notch signaling and elevated Akap12 expression disrupt the ability of renal epithelial cells to form spherical structures with a single lumen when grown embedded in matrix. We conclude that Notch signaling regulates *Akap12* expression to ensure normal primary cilia length and renal epithelial morphogenesis, and suggest that diseases associated with defective Notch signaling, such as Alagille syndrome, maybe mechanistically related to ciliopathies.

**Translational Statement:** The current study examines how a reduction in Notch signaling results in abnormal renal tubule formation, as occurs in Alagille Syndrome patients with mutations in *JAG1* or *NOTCH2*. The finding that reduced Notch signaling results in abnormally long cilia is suggestive that some of the clinical manifestations in Alagille Syndrome, such as small cystic kidneys, may originate due to defective cilia function. Linking Notch to primary cilia also opens up the possibility that coinheritance of mutations in ciliopathy genes along with a mutation in *JAG1* or *NOTCH2* may enhance the severity of the clinical phenotypes such as cystic kidney disease and may explain the variable occurrence and onset of kidney disease among Alagille Syndrome patients.

## Introduction

Epithelial tubule formation, growth and patterning are tightly regulated processes that occur during the normal development of multiple organs including the salivary gland, mammary gland, lungs and kidneys. Defective formation, growth and/or maintenance of tubular epithelial structures can severely impact the functions of organs and result in diseases such as cystic kidney disease and renal cell carcinoma.^1–3^ Mutations in one or more of a large set of genes results in cystic kidneys in which segments of nephrons and/or collecting ducts, composed of tubular epithelia, are abnormally enlarged and can progressively disrupt overall renal structure and function. Many of the cystic kidney disease genes have been linked to primary cilia and/or centrosomes, which may regulate epithelial cell division among other processes to ensure normal tubule formation and growth.^4–7^

Patients with Alagille syndrome carry loss of function mutations in *NOTCH2* or *JAG1* and variably develop small multicystic kidneys.^8–11^ Additionally, gain-of-function mutations in *NOTCH2* are associated with polycystic kidneys that occur in Hajdu-Cheney syndrome (OMIM10250) and serpentine fibula polycystic kidney syndrome (SFPKS; OMIM600330) patients among other phenotypes. Conditional inactivation of *Notch1* and *Notch2* genes within the nephrogenic lineage results in defective suppression of Six2^12^ and subsequently impaired conversion of renal vesicles into S-shaped bodies in mice.^12,13^ Inactivation of *Notch1* and/or *Notch2* floxed alleles or *RBPJ* floxed alleles from the cap mesenchyme stage onwards resulted in defective nephrogenesis with the variable occurrence of multicystic kidneys of proximal tubule origin.^14,15^ The occurrence of renal cysts correlated with the degree of Notch signaling inactivation, with penetrance reaching a 100% when both alleles of *Notch1* and *Notch2* were inactivated in the cap mesenchyme.^14^ Together these observations in mice and humans confirm a role for Notch signaling in renal tubule morphogenesis, but the mechanisms by which Notch signaling suppresses renal tubular cysts remain unclear. Primary cilia and basal bodies that anchor cilia are linked to cystic kidney diseases, but whether Notch signaling requires primary cilia ^16,17^, or is functionally linked to basal bodies/centrosomes during renal tubule morphogenesis remains to be determined. In order to understand the involvement of Notch signaling in the regulation of renal tubule morphogenesis and in the etiology of kidney cysts we investigated the functions of Notch signaling in renal epithelia and have determined that it ensures normal primary cilia length and structure.

## Results

### Notch signaling regulates tubule morphogenesis even after S-shaped stage of nephrogenesis to prevent cyst formation

To better define the stages at which loss of Notch signaling allows for renal tubular cyst formation we suppressed Notch signaling mediated transcription after the formation S-shaped bodies (SSB) within nephron forming cells and the developing collecting duct cells. We hypothesized that since Notch1 and Notch2 are expressed in SSBs,^18^ Notch signaling may regulate a cellular process during or after the SSB stage whose absence promotes cyst formation. To test this idea transgenic mice with *Pax8* regulatory region driving reverse tetracycline transactivator (rtTA)^19^ were bred with mice carrying a tet-response-element (TRE) driving dominant-negative mastermind-like (dnMaml), a stoichiometric inhibitor of Notch/RBPJ-mediated transcription.^20^ *Pax8->rtTA* drives the expression of rtTA in the nephrogenic lineage from the SSB stage onwards and in collecting ducts of the developing kidney.^19^ In these *Pax8->rtTA;TRE->dnMamL* mice, expression of dnMamL was induced by providing doxycycline (dox) in the drinking water. Dox treatment throughout the pregnancy of *TRE->dnMaml* females time mated with *Pax8->rtTA* male mice and analysis of kidneys of new born pups revealed cortical cysts (arrowheads Fig.1B) that are LTL+ (arrowheads in Fig.1D) and hence of proximal tubular origin only in *Pax8->rtTA;TRE->dnMamL* mice (Fig.1B&D) and not in dox treated control littermates (Fig.1A&C). The presence of enlarged proximal tubular segments (boxed area in Fig.1D) is suggestive that Notch signaling has a role in this nephrogenic lineage after SSB stage, and in the absence of Notch signaling proximal tubular cysts can form. Additionally, medullary cysts are also present in *Pax8->rtTA; TRE->dnMamL* mouse kidneys (asterisks in lumen of enlarged ducts in Fig.1B) but not in control littermate kidneys (Fig.1A&C). The expression of dnMaml in the developing collecting ducts, which are derived from a different progenitor, results in collecting duct cysts lined with Aqp2+ epithelia (Fig.1D).

**Figure 1.**
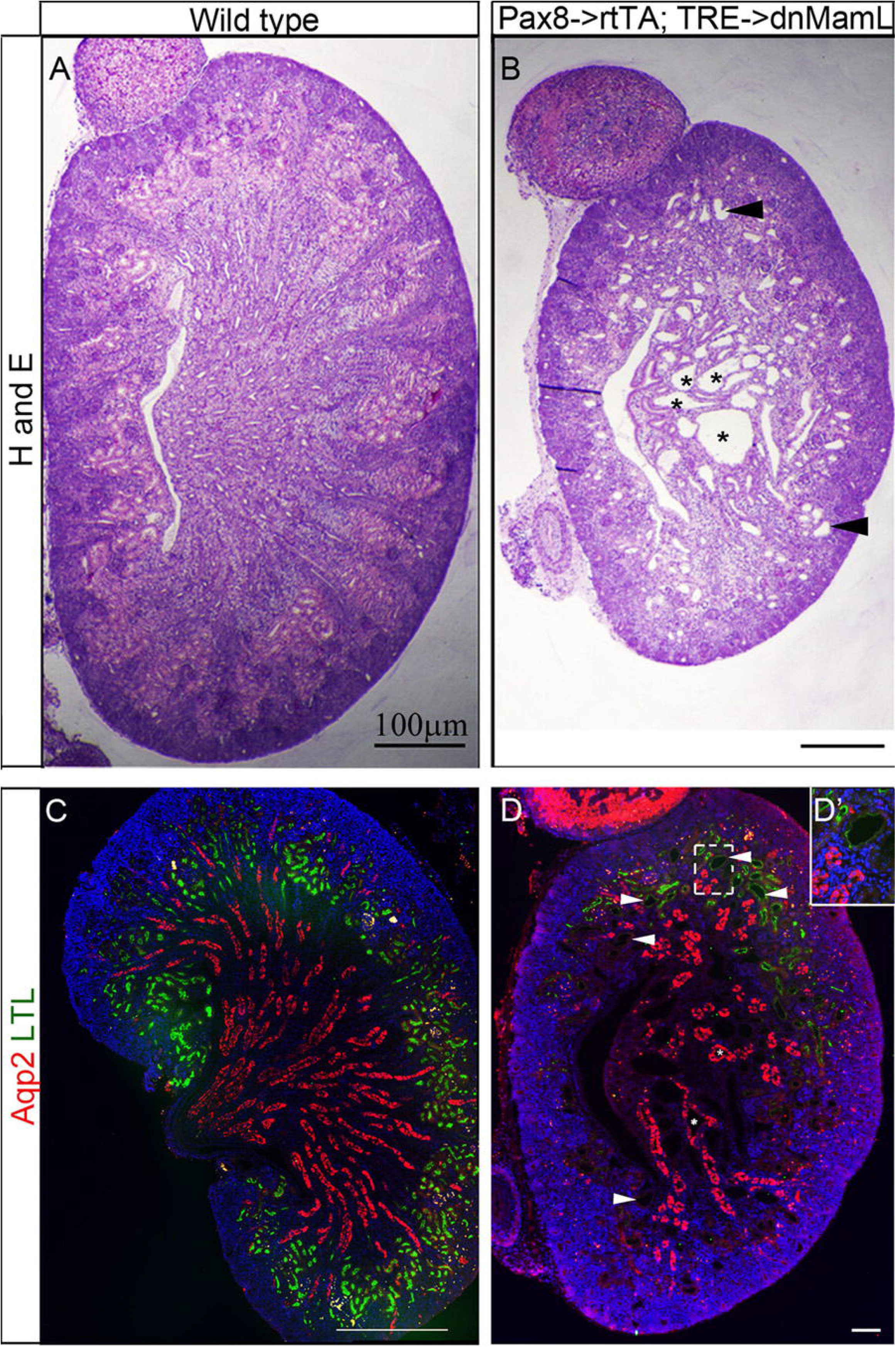
Notch signaling is required within nascent nephrogenic structures even after the s-shaped stage during nephrogenesis for normal proximal tubule morphogenesis and cyst suppression. A & B are hematoxylin & eosin stained kidney sections from post-natal day zero mice. Kidney sections from wild type mice (A, C) appear to be normal, while kidney sections with Notch-signaling inhibition (B, D) are smaller and show the presence of cortical cysts (arrowheads in B and D) and medullary cysts (asterisks in lumen of cysts in B and D). The cortical cysts are of proximal tubule origin as they stain positive for LTL a marker of proximal tubules (arrowheads in D), while the medullary cysts are Aqp2-positive and hence are of collecting duct origin (asterisks in D). D’ is a higher magnification view of the boxed area in D, which contains a proximal tubule cyst.

To determine if there is a critical window during kidney development during which Notch activity prevents formation of renal tubular cysts, we provided dox for durations of two days during embryogenesis and analyzed the kidneys at birth. Smaller kidneys with multiple cysts developed in *Pax8->rtTA;TRE->dnMamL* mice analyzed at birth when pregnant dams were given dox between days E12.5 to E14.5, E14.5 to E16.5 or E16.5 to E18.5 (Fig. 2B,D&F) while control littermates developed normal kidneys (Fig. 2A,C&E). These results indicate that Notch signaling is required from E12.5 to E18.5 to ensure normal renal morphogenesis and to prevent proximal tubular and collecting duct cysts. Additionally, these observations reveal that even acute inhibition of Notch signaling is sufficient to disrupt tubule morphogenesis leading to renal cyst formation during this window.

**Figure 2.**
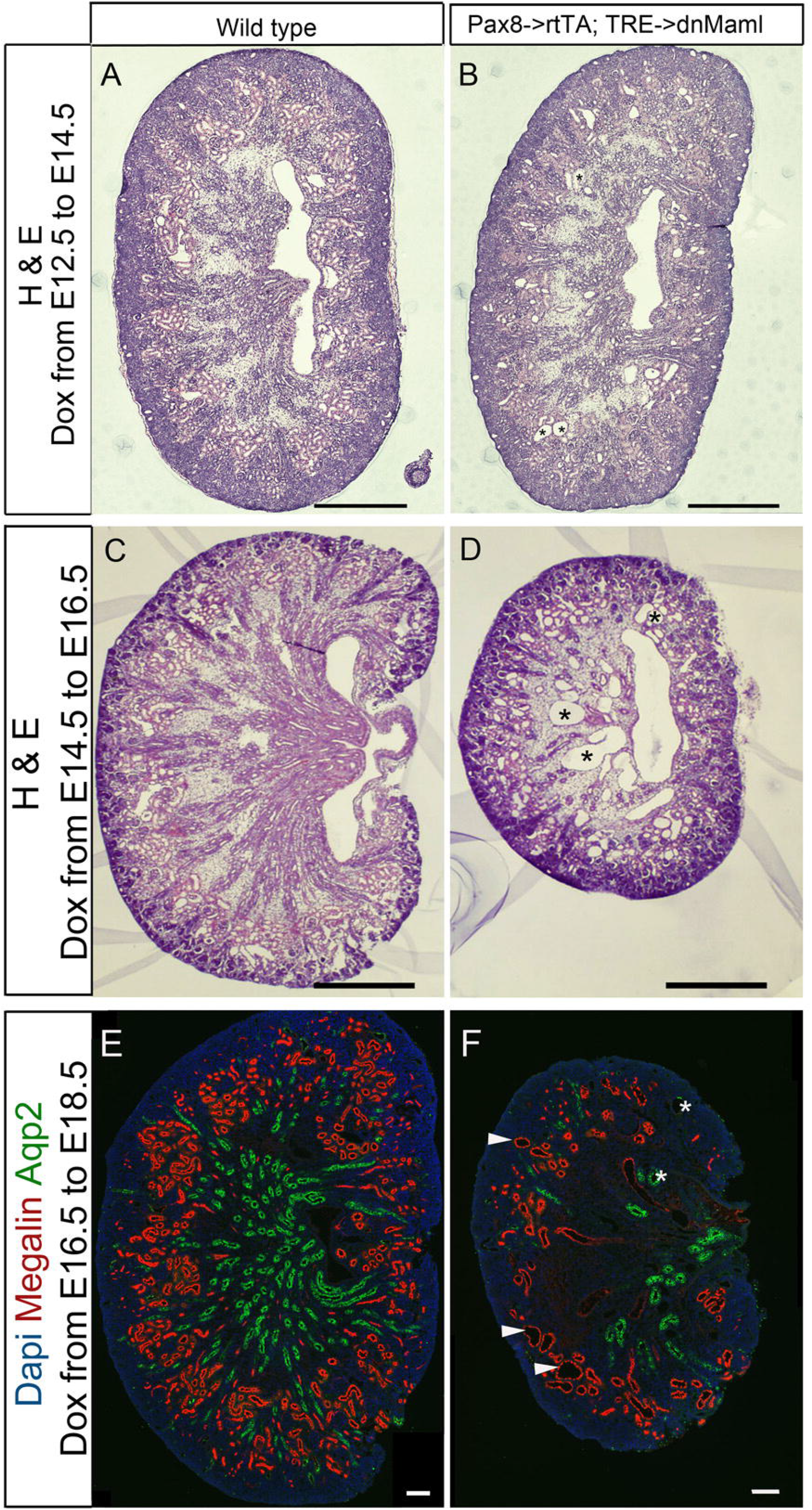
Acute inhibition of Notch signaling at any point during nephrogenesis is sufficient to disrupt proper tubule formation and allows for the formation of renal tubular cysts. **A-D** Hematoxylin & eosin stained kidneys with uninterrupted Notch-signaling (*WT, Pax8->rtTA, or TRE->dnMamL mice*) or mutant littermates. Wild type kidneys (A, C) show no cysts while their Notch-inhibited littermates (B, D) show presence of cysts at all time points tested. Asterisks in B mark small cysts in mutant kidneys, while the asterisks in C mark large prominent cysts. **E,F** Immunohistochemistry of wild type (E) and Notch-signaling-inhibited (F) kidneys show that cysts are of proximal tubule and collecting duct origin, as evidenced by staining with a proximal tubule marker Megalin (arrowheads in F) and a collecting duct maker Aqp2 (asterisks in F). Scale bars are 500μm.

### Inhibition of Notch signaling in renal epithelial cell cultures results in abnormal spheroid formation

To begin determining the cellular processes and genes regulated by Notch signaling within renal epithelia we established stable MDCK cell lines expressing dnMaml fused with EGFP at the c-terminus (Fig.3A-C). Compared with EGFP expression, which localizes throughout the cells (Fig.3A), dnMaml::EGFP predominantly localizes within nuclei (Fig. 3B). The Notch responsive TP-1-Luciferase reporter activation by Notch1 intracellular domain (NICD) was inhibited in MDCK colonies expressing dnMaml::EGFP compared with MDCK colonies expressing EGFP (Fig. 3C). We next compared the ability of parental MDCK cells and two of the dnMaml::EGFP expressing MDCK cell lines for their ability to form spheroids when grown embedded in collagen matrix. The ability of epithelial cells to form spherical epithelial structures with a single lumen is in part dependent on regulated orientation of cell division,^21,22^ epithelial polarization and other cellular processes. Interestingly, the ability of MDCK cells to form a sphere with a single lumen is severely compromised with expression of dnMamL (Fig. 3D-H). The formation of multiple lumens by dnMaml expressing epithelia is consistent with the abnormal orientation of renal epithelial cell division previously observed within the proximal tubules of mice with Notch-signaling deficient kidneys.^14^

**Figure 3.**
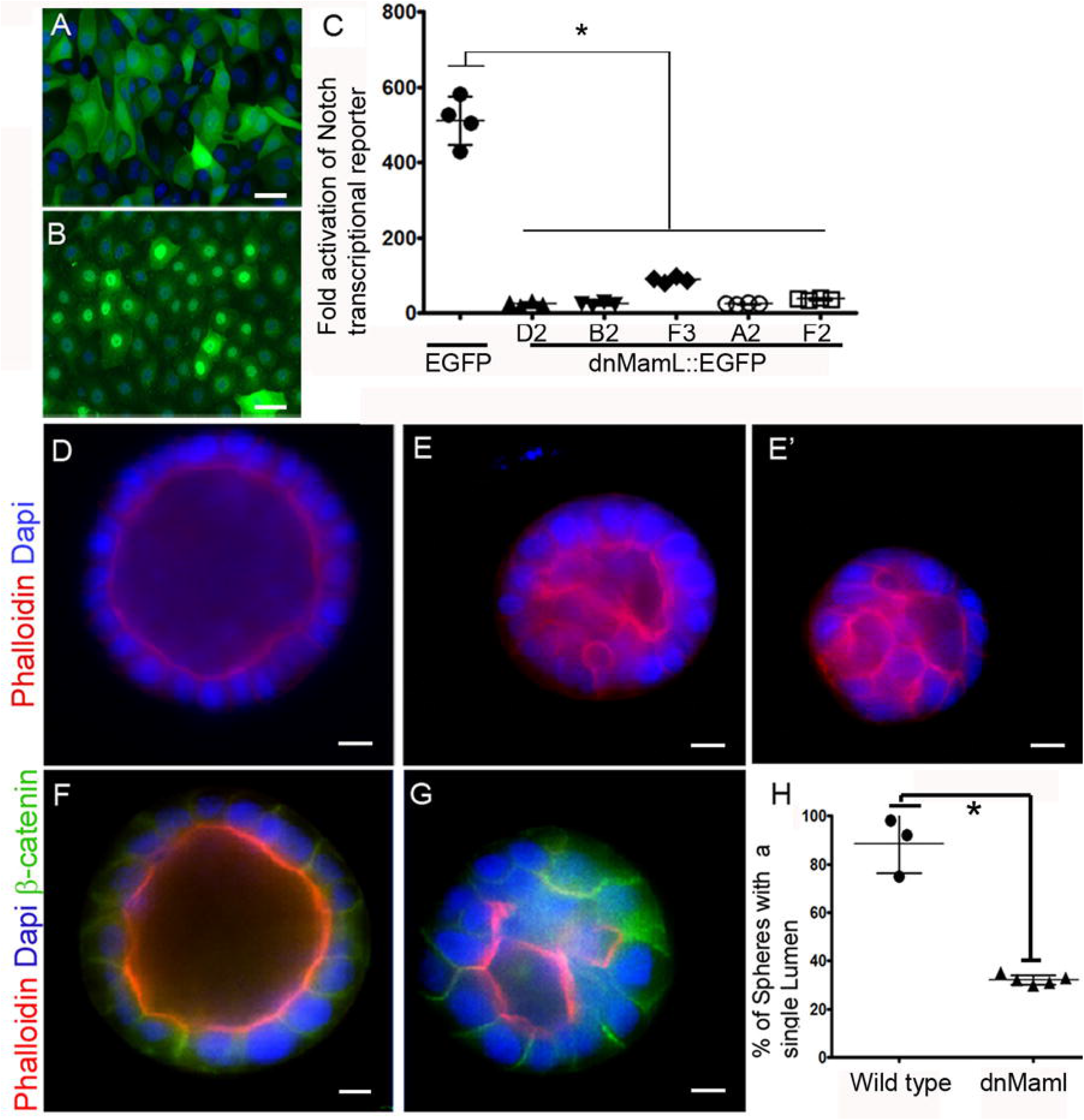
Stable expression of dominant-negative Mastermind-like in MDCK cells inhibits Notch signaling and impairs their ability to form spheroids with single lumens. A, B Fluorescent microscopy shows homogeneous GFP+ population indicative of EGFP or dnMaml::EGFP expression. In cells transfected with EGFP (A), fluorescence is localized predominantly in the cytosol, while in cells expressing dnMamL fused with EGFP, fluorescence localizes to the nucleus. C. Cells expressing dnMaml::EGFP show inhibition of Notch-signaling mediated transcriptional activation as observed by reporter assays. Stable clones expressing either EGFP alone or dnMamL::EGFP were tested for their ability to activate the Notch-responsive TP-1 promoter. Compared with an EGFP-expressing clone that showed robust activation of TP1-1 promoter in response to transient expression of activated Notch1 (NICD) relative to empty vector, the ability of NICD to activate transcription is significantly reduced in clones with dnMaml::EGFP expression. Asterisk denotes p<0.05 when comparing each dnMaml expressing clone with EGFP expressing clone as determined by student t-test, n=4 per clone. D-H. MDCK cells (D, F) form 3D spheroids with single lumen and show proper cell-cell junctions as seen by β-catenin staining for adherens junctions on the basolateral membrane. Phalloidin stains filamentous actin to illuminate epithelial cell structure. Cells with inhibition of Notch signaling (E, E’, G) form visually smaller spheroids with multiple lumens. H. Quantification of spheroid formation assay reveals that on average 88% of the spheroids formed by wild type cells show a single lumen (n=100, three repeats), while only 30% of cells with Notch-signaling-deficiency form spheroids with single lumen (n=100, five repeats using three different dnMaml::EGFP expressing MDCK cell lines). Asterisk denotes p<0.05 student t-test. Scale bars are 10μm.

### Inhibition of Notch signaling results in longer primary cilia with abnormal ciliary microtubule arrangements

Numerous genetic mutations that result in cystic kidney diseases have been associated with defective cilia and basal bodies/centrosomes.^23,24^ To determine the underlying cause of abnormal tubule morphogenesis in Notch signaling deficient kidneys, we analyzed primary cilia and centrosomes. MDCK cells (n=85) formed cilia with an average length of 1.3+/- 0.6μm (Fig. 4A), whereas two independent dnMaml expressing cell lines (n=100 and n= 60) had a combined average cilia length of 3.26+/- 1.8μm (Fig.4B &C; p<0.05, test-test comparing parental MDCK cells versus dnMaml clones 1 and 2 combined). We similarly observed increased cilia length in serum starved NIH3T3 cells following DAPT treatment to block Notch signaling compared with DMSO treated NIH3T3 cells (Supplementary Fig. S1).

**Figure 4.**
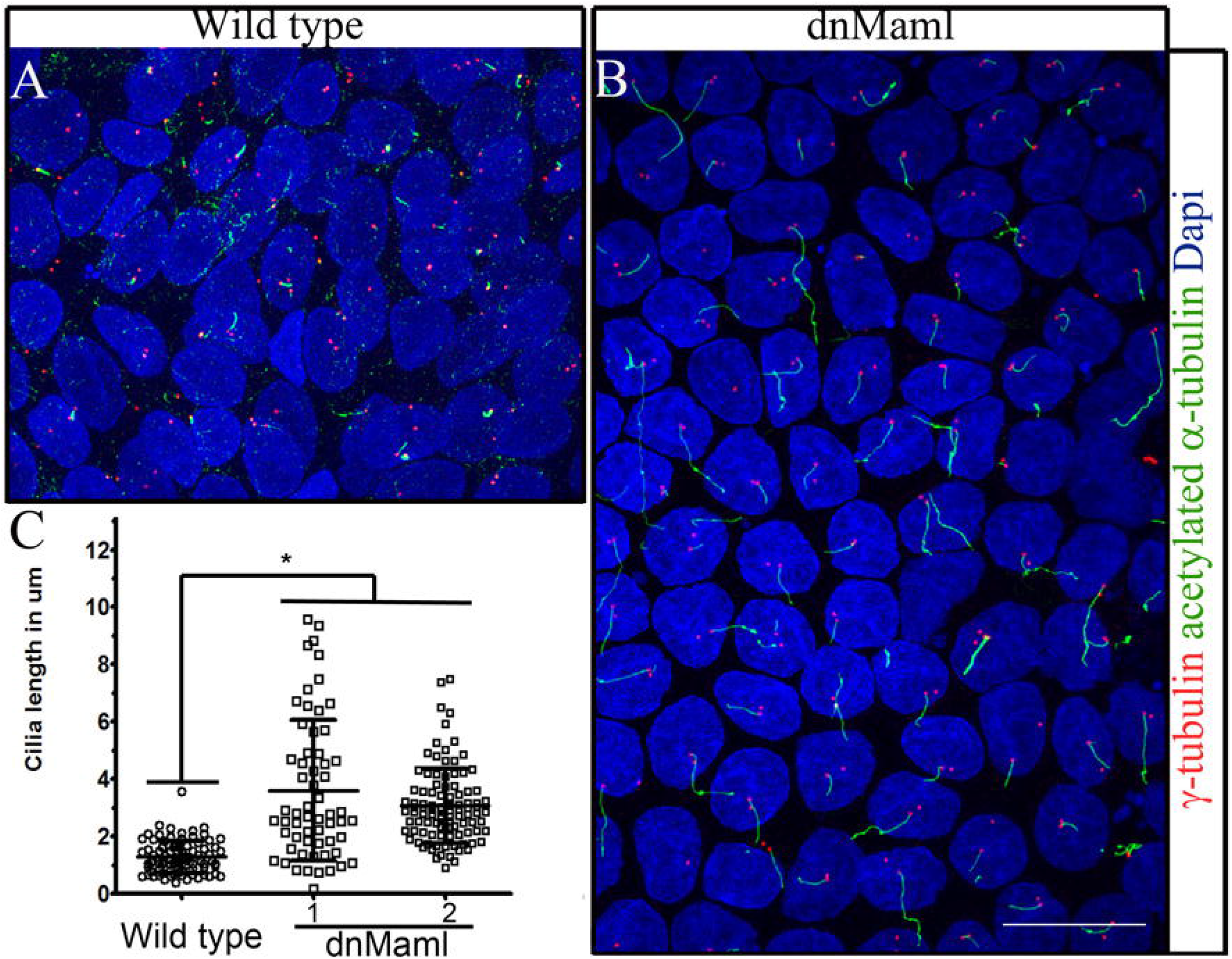
Inhibition of Notch-signaling in cultured renal epithelial cells results in abnormally long cilia. Wild type (A) and Notch-signaling-deficient (B) MDCK cells grown on transwell membrane were induced to form cilia by a combination of cell-cell contact inhibition, and serum starvation. Cilia were visualized by staining for acetylated α-tubulin and centrosomes were labeled with γ-tubulin. C. Cilia measurements conducted in wild type MDCK versus two independent MDCK cell lines with dnMamL-EGFP expression revealed significantly longer cilia in Notch-signaling-deficient cells. Asterisk denotes p<0.05, following t-test comparing wild type (n=85), versus two Notch signaling inhibited cell lines (n=160).

This increase in primary cilia length also occurs in mouse kidneys with conditional inactivation of Notch signaling (Fig. 5). Scanning EM revealed cysts and longer primary cilia in three week old *Rarb2-Cre*; *RBPJ^f/f^* mouse kidneys (Fig. 5B, D and F) when compared with control littermate kidneys that had no cysts and shorter cilia (Fig. 5A, C and E). To quantify the increase in cilia length we stained Notch signaling deficient kidneys with an antibody against acetylated-tubulin and megalin to mark proximal tubules (Supplementary Fig. S2). We observed that on average the cilia length in control littermates was 4.22+/-1.08μm versus 9.04 +/-0.57 μm (Supplementary Fig. S2).

**Figure 5.**
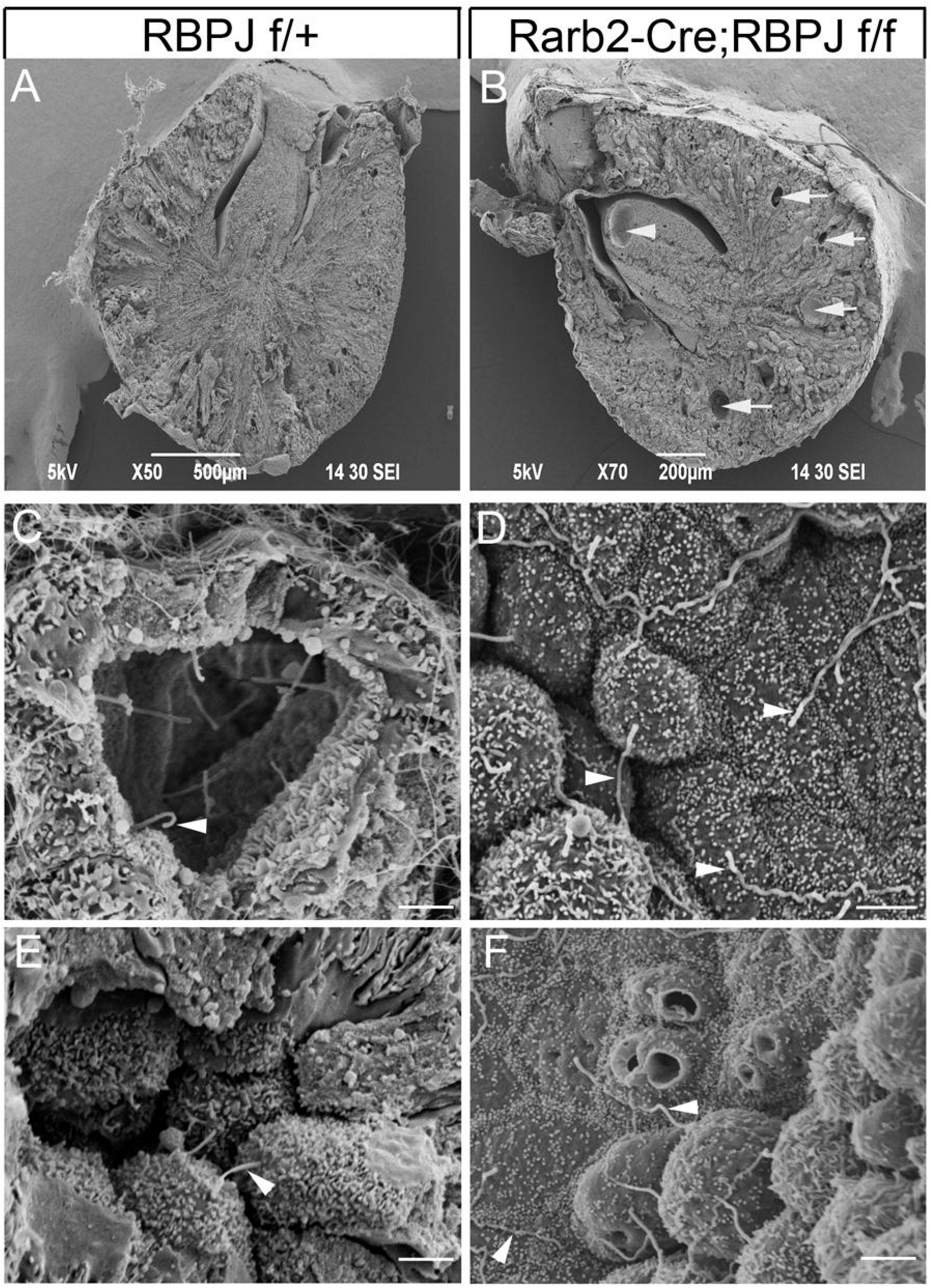
Mouse kidneys with conditional inactivation of Notch-signaling have abnormally long renal epithelial cilia. A-F Scanning electron micrographs of three-week-old wild type (A, C, E) and Rarb2-Cre; RBPJ^F/F^ (B, D, F) mouse kidneys. Kidneys with conditional inactivation of Notch-signaling show the presence of cysts (arrowheads in B) whereas kidneys of control littermates (A) show no cysts. Higher magnification images show the presence of primary cilia of normal length in wild type kidneys (arrowheads in C & E) while the cilia seen in mutant kidneys are significantly longer (arrowheads in D & F). Scale bars in C-F are 2μm.

We next examined the structure of the cilia in wild type and dnMaml expressing MDCK cells, as well as in wild type and Notch-signaling-deficient mouse kidneys. Consistent with primary cilia being present on the apical surface of quiescent renal epithelia we observed a 9+0 microtubule arrangement within the ciliary axoneme of wild type renal epithelia in moue kidneys (Fig. 6A) and MDCK cells (data not shown). In contrast, we observed that a majority (~85%) of cilia analyzed in Notch-signaling-deficient renal epithelia, including dnMaml expressing MDCK cells (Fig.6 G&H) and mouse kidneys deficient in Notch signaling (data not shown) deviated from the 9+0 microtubule arrangement within the ciliary axoneme. The abnormal arrangements included 6 to 8 outer pairs of microtubules plus one central pair (Fig. 6 G&H). Additionally, there is an electron dense sheath around the central pair and electron dense structures at locations similar to where outer dynein arms are normally present in motile cilia (Fig. 6G&H). Only a minority (~15%) of the cilia observed in Notch-signaling-deficient renal epithelia had normal 9+0 microtubule arrangement, although many of these also had electron dense regions resembling outer dynein arms. The transition zone consisting of a necklace like structure and basal body consisting of triplet microtubules appeared normal in both wild type and Notch-signaling-deficient renal epithelia (Fig. 6 B, C, E &F). Overall, the cilia are longer in Notch-signaling-deficient renal epithelia and the structure appears to deviate from the normal 9+0 microtubule organization found within the axoneme of primary cilia.

**Figure 6.**
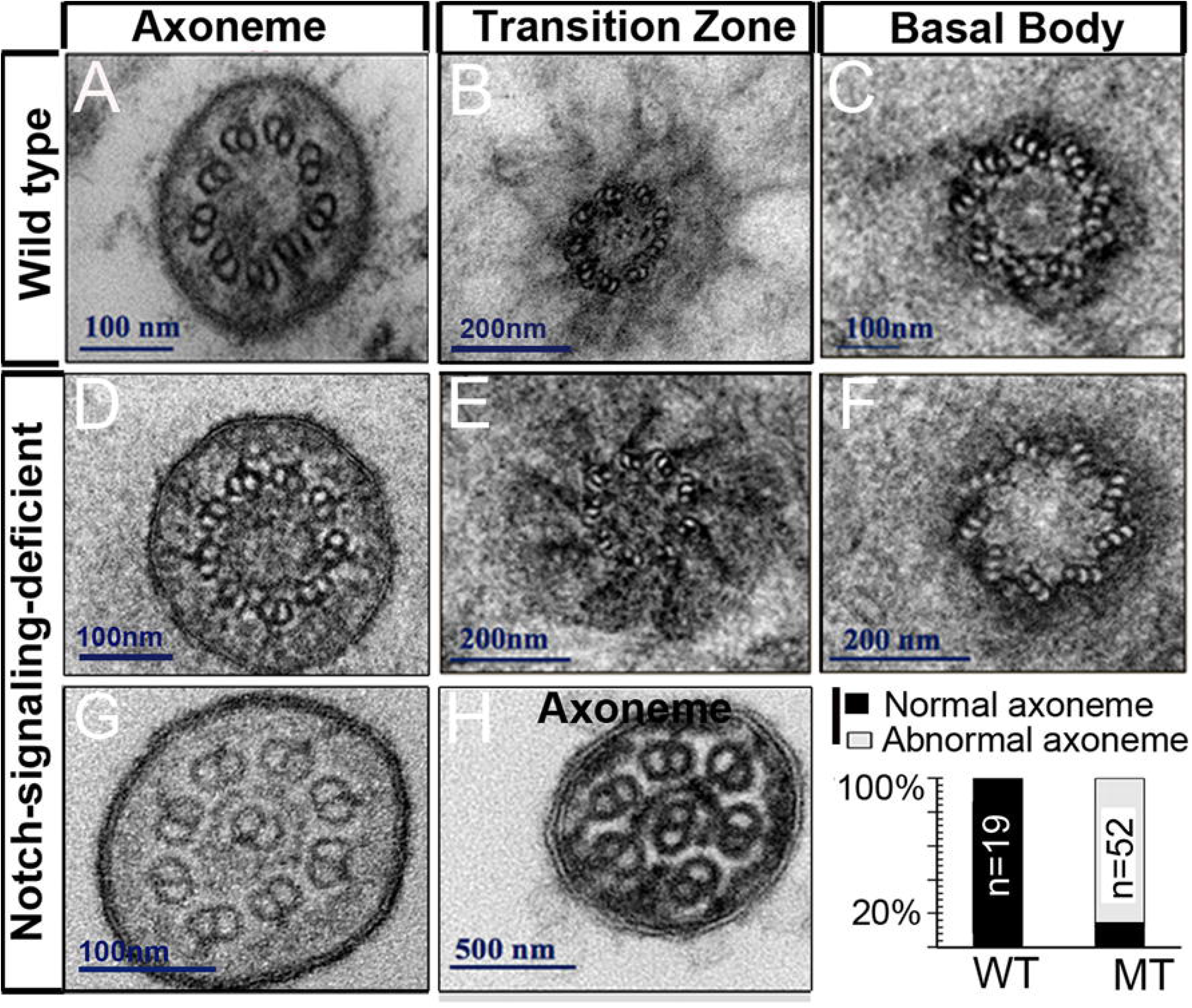
Renal epithelial cells with Notch-signaling-deficiency show abnormal ciliary ultrastructure by TEM. Wild-type renal epithelial cilia (A, B, C) show normal 9+0 arrangement of microtubules at the axoneme, transition zone and basal body (n=19). Only a minority (~15% of 52) of Notch-signaling-deficient cilia showed normal 9+0 microtubule arrangement within the axoneme (D). Defects seen in the Notch-signaling-deficient ciliary axoneme include 8 outer pairs (G) or 6 outer pairs (H) along with a central pair, and a sheath of electron dense material around the central pair (H). I. Quantification of normal versus abnormal ciliary ultrastructure by TEM reveals that a majority of the cilia in the Notch-signaling-deficient cells have abnormal axonemes.

### *Akap12* expression level increases with the loss of renal epithelial Notch signaling

The canonical Notch pathway components such as Notch1, Notch2, and RBPJ are required to suppress renal cysts and regulate gene expression, which dnMaml can inhibit. This led us to hypothesize that Notch signaling regulates genes that suppress renal cysts by governing cilia length and/or orientation of cell division. To identify Notch regulated cyst suppressor genes we compared the gene expression profiles of quiescent polarized MDCK cells with and without dnMaml expression by microarray analysis (data not shown). From the list of differentially expressed genes we focused on genes associated with centriole/centrosome, cilia and/or cystic kidney disease and validated the increased expression of *Ahi1, Akap12, Bicc1, Ift172*, and *Nphp1* by RT-qPCR in MDCK cells with dnMaml expression (Fig. 7A). For further validation we compared the gene expression in kidneys of E18.5 *Pax8->rtTA;TRE->dnMamL* versus wild-type embryos that had been exposed to dox from E16.5, since this results in dnMaml-dependent renal cysts at birth (Fig. 2). By RT-qPCR we determined that *Ahi1* and *Akap12* are up regulated in dnMaml expressing E18.5 kidneys, with only *Akap12* being statistically significant, while the other differentially expressed genes of interest identified in MDCK cells were not significantly up-regulated following dnMaml expression *in vivo* (Fig. 7B).

**Figure 7.**
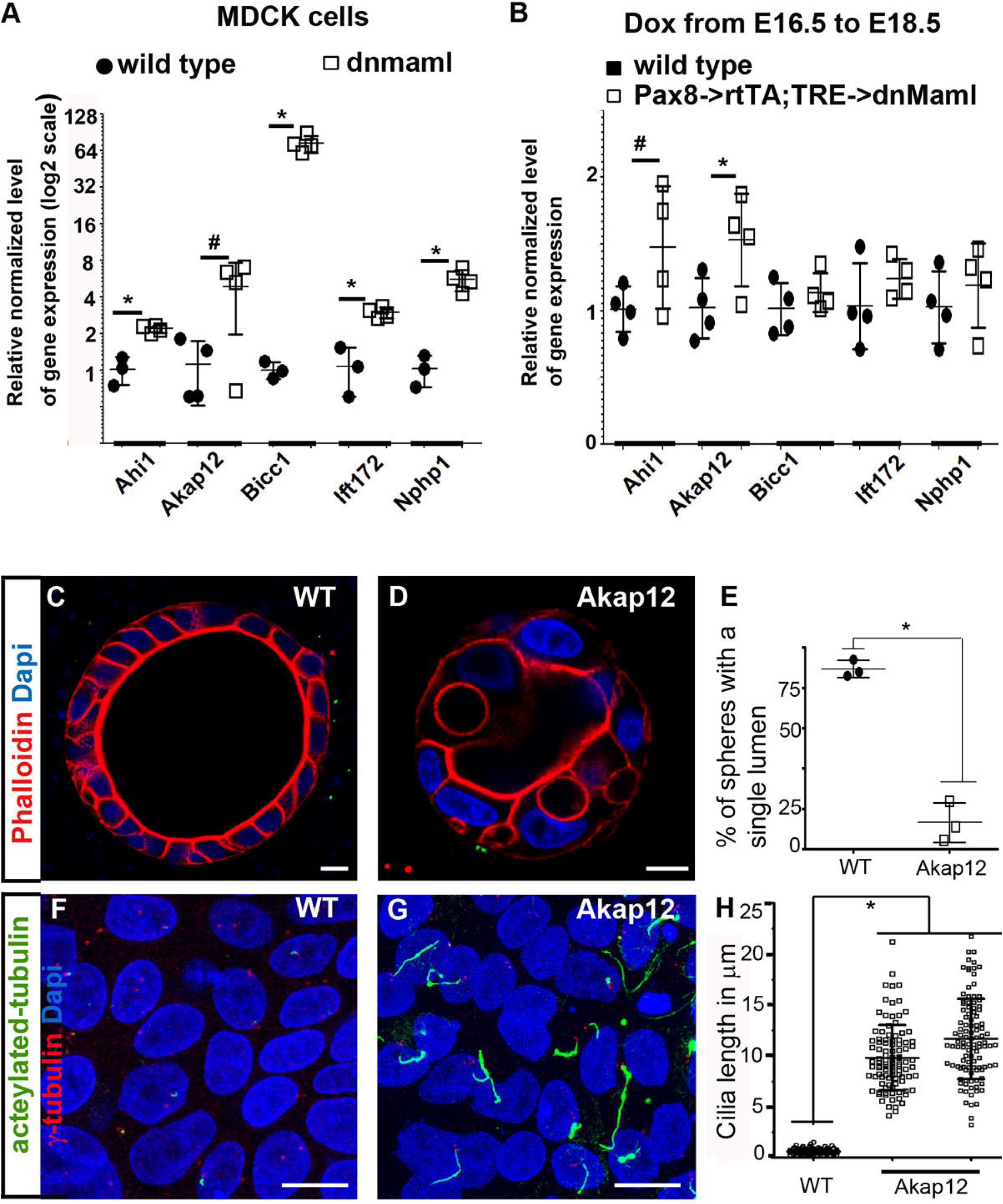
Ectopic expression of Akap12 recapitulates the ciliary length and spheroid formation defects seen in Notch-signaling-deficient renal epithelia. A. RT-qPCR validation of genes related to cilia, centrosomes, and cell cycle regulation identified by microarray analysis that were differentially expressed in Notch-deficient renal epithelial cells compared with wild-type cells. Asterisks denote p<0.05, n=3 or 4 for each condition, # denotes p=0.078 n=4 per condition as determined by t-test. B. RT-qPCR shows significant upregulation of *Akap12* in E18.5 kidneys of *Pax8->rtTA; TRE->dnMamL* mice compared to wild type littermates given Dox from E16.5. Asterisk denotes p<0.05, n=4 per group. Although, *Ahi1* expression also increases this was not statistically significantly different between genotypes, # denotes p=0.1, n=4 per group as determined by t-test. C-E. MDCK cells with ectopic overexpression of Akap12 form spheroids with multiple lumens (D) compared to wild type cells (C). E. On average 89% of parental cells form spheroids with a single lumen (n>50 in each of three replicates), while on average 13% of Akap12 overexpressing cells formed spheroids with a single lumen (n>45, three separate clones with Akap12 overexpression). * denotes p<0.05, student t-test. F-H. Parental MDCK cells (F) and those with ectopic Akap12-epression (G) were grown on transwell membrane and induced to form cilia by a combination of cell-cell contact inhibition, and serum starvation. Cilia were visualized by staining for acetylated α-tubulin and centrosomes were labeled with γ-tubulin. H. Cilia measurements conducted in wild type MDCK versus two independent MDCK cell lines with ectopic stable Akap12 expression revealed significantly longer cilia in cells with ectopic Akap12 expression. Asterisk denotes p<0.05, when comparing wild type (n=120) versus Akap12 expressing (n=207) cells by t-test. Scale bars are 10μm.

### Ectopic expression of Akap12 triggers an increase in cilia length and alters the ability to form spheroids with a single lumen

Previous studies report that the scaffolding protein A-kinase anchoring protein 12 (Akap12) forms a complex with mitotic kinases, Aurora A and Plk1 to regulate mitotic spindle orientation during cell division in seminiferous tubules.^25^ Considering that Akap12 is up regulated in kidneys with a brief inhibition of Notch signaling, and is known to regulate orientation of cell division, we tested whether increased expression of Akap12 could explain the abnormal cellular phenotypes observed in Notch-signaling-deficient renal epithelia. Stable expression of Akap12 in MDCK cells resulted in abnormal spheroid formation (Fig. 7C-E). While a majority (89%) of parental MDCK cells formed spheroids with a single lumen when grown in 3D collagen matrix (Fig. 7C), only 13% of the cells with ectopic expression of Akap12 formed spheroids with a single lumen (Fig. 7D). Additionally, when MDCK cells with ectopic Akap12 expression were grown to confluence on transwell membranes they formed abnormally long primary cilia with an average length of 10.8+/- 3.7μm (n=207 from two Akap12 expressing clones) compared to the parental cells with an average cilia length of 0.65+/- 0.24μm (n=120) (Fig. 7F-H). These observations are suggestive that Notch signaling mediated repression of *Akap12* in renal epithelia is part of the mechanism by which Notch signaling ensures normal renal primary cilia length and tubule morphogenesis.

## Discussion

### Alagille Syndrome may in part be a ciliopathy

Alagille syndrome patients are observed to have abnormalities in multiple organs including the kidney, liver and heart, and are differentially diagnosed from ciliopathies based on genetic testing. Hepatorenal fibrocystic diseases such as autosomal dominant and recessive polycystic kidney diseases (ADPKD and ARPKD), Meckel syndrome, and nephronophthisis (NPHP) with liver disease have been referred to as ciliopathies, based on the observation that proteins defective in these diseases localize or are associated with primary cilia.^26,27^ In common with Alagille Syndrome, ciliopathies are associated with kidney, liver and heart disease. There is at least one case report of a patient with kidney, heart and liver abnormalities who was initially considered to have Alagille Syndrome who was then identified to have biallelic mutations in *NEK8*, also known as *NPHP9*, a ciliopathy associated gene.^28^ Our observation that Notch signaling deficiency results in longer cilia, which also occurs in renal epithelia with mutations in *polycystin1* (PKD1),^29^ is suggestive that Notch signaling deficiency should also be considered a ciliopathy.

### The longer cilia in Notch-signaling-deficient renal epithelia maybe related to the role of Notch signaling in inhibiting the multi-ciliated cell fate selection

Considering that inhibition of Notch signaling promotes multi-ciliated cell fate in organs such as the lung among others in mice,^30^ in the frog skin,^31^ and in the pronephros of zebrafish,^32^ it is possible that Notch signals to suppress the multi-ciliated cell fate which maybe abnormally turned on with the loss of Notch signaling in renal epithelia. Multi-ciliated cells have motile cilia, which have a distinct substructure from primary cilia in that they are composed of nine outer pairs of microtubules plus an inner central pair, along with dynein arms. We observed that cilia on Notch-signaling-deficient renal epithelia frequently have a centrally located pair of microtubules, as a well as structures that resemble a central sheathlike structure and outer dynein arms (Fig. 6). Additionally, the multi-ciliated program involves centriole amplification required to anchor the numerous motile cilia. Interestingly, we occasionally observed multiple centrioles in the renal epithelia lining the cysts of Notch-signaling-deficient kidneys (Supplementary Fig. S3).

### Notch signaling is required for normal tubule morphogenesis within nephron forming structures from after the s-shaped stage, and is required for normal morphogenesis of the collecting ducts

Previous studies have identified roles for Notch signaling in the kidney, including the suppression of Six2 within the nephrogenic lineage^12^ as well as the subsequent conversion of renal vesicles (RV) into S-shaped bodies (SSBs).^13,33,34^ These early requirements of Notch signaling may be responsible for the suppression of kidney proximal tubular cysts,^14,15^ or alternatively there may be a function for Notch after SSB formation. Here we initiated the inhibition of Notch signaling from the SSB stage onwards within the nephron forming cells. This resulted in kidney proximal tubular cysts revealing a function for Notch signaling in tubule morphogenesis from after the formation of s-shaped structures. Additionally, we observed that two-day windows of dox induced dnMaml expression between E12.5 to E18.5 resulted in renal cysts. Since nephrogenesis is repeatedly initiated from E12.5 onwards, it appears that all nephrons are susceptible to loss of Notch-mediated cyst formation, and even transient inhibition of Notch signaling alters tubule morphogenesis. Apart from the nephrons, collecting duct morphogenesis is also altered resulting in collecting duct cysts. Interestingly, dox treatment from E14.5 to E16.5 resulted in the most severe medullary collecting duct cysts and smaller kidneys compared with treatments from E12.5 to E14.5, as well as from E16.5 to E18.5. Although, collecting duct branching occurs from E12.5 through E18.5 there appear to be critical periods during duct morphogenesis that are highly sensitive to loss of Notch signaling.

### Akap12 is a gene regulated by Notch signaling in renal epithelia that is also involved in regulating tubule morphogenesis

A function of Akap12, also known as Gravin or SSeCKS, is to recruit Aurora A and Polo-like kinase 1 (Plk1) to the mother centriole.^25^ The restricted localization of Aurora A and Plk1 to the mother centriole allows for symmetric orientation of cell division.^25^ Numerous studies have observed a role for Aurora A in mitotic spindle orientation in epithelial cells.^35–38^ We observe that loss of Notch signaling increases *Akap12* expression, and increased ectopic Akap12 results in abnormal renal tubule morphogenesis. Consistent with increased levels of Akap12 resulting in epithelial abnormalities, in addition to mutations, amplifications of *AKAP12* have been associated with prostate and ovarian cancers.^39,40^

## Methods

### Mice

All experiments involving mice were approved by the Sanford Research IACUC. Mice were genotyped following a universal PCR genotyping protocol.^41^ Primer sequences are available upon request. Wild type control groups consisted of age and where possible sex matched littermates housed along with mice with conditional inactivation of Notch signaling. Doxycycline (1mg/mL; Sigma-Aldrich) was administered in drinking water containing 5% sucrose to adult mice during which they continued to be fed normal chow ad libitum. *Pax8->rtTA;TRE->dnMaml* male mice were time mated with C57Bl6/J female mice. Doxycyline was provided to females either from date of observed plug or for two days during pregnancy between embryonic day (E) 12.5 to 18.5. Post-natal day 0 mouse kidneys were isolated for histology or RNA extraction. Offspring from mating *Rarb2-Cre;RBPJ ^+/f^* male mice with *RBPJ^f/f^* females were euthanized for analysis of kidneys by scanning EM and for primary cilia length determination. Offspring from *Fgf20-Cre;Notch1^+/f^;Notch2^+/f^* bred with *Notch1^f/f^;Notch2^f/f^* were euthanized for analysis of kidney by TEM. Offspring from *Six2-Cre;Notch2^f/f^* bred with *Notch1^f/f^;Notch2^f/f^;Cent2-GFP* were euthanized for determining centriole number per epithelial cell within their kidneys. Kidneys were fixed in OCT, sectioned and stained with phalloidin to visualize actin filaments and GFP fused centrioles by confocal microscopy.

### Histology and immunohistochemistry

Kidneys were fixed in 4% PFA or Bouin’s fixative overnight at 4°C, washed in 70% ethanol, paraffin embedded and sectioned at 12μm thickness. Prior to immunostaining, the sections were de-paraffinized in xylene, rehydrated and, boiled for 20 min in Trilogy (Cell Marque) for antigen unmasking. Sections were blocked in PBS containing 1% bovine serum albumin (BSA), 0.2% powdered skim milk, and 0.3% Triton X-100 for at least 15 min at RT prior to incubation with primary antibodies overnight. For direct visualization of GFP or tdtomato the mouse kidneys were fixed in 4% PFA for 1 hour, rinsed in PBS, and incubated in 15% sucrose for 24 hours and then another 24 hours in 30% sucrose prior to embedding in OCT and sectioning at 12μm thickness. The primary antibodies used in this study include: Aqp2 (1:200, sc-9882, Santa Cruz Biotechnology), Megalin (1:200, ab184676, Abcam), and acetylated-α-tubulin (1:500, T7451, Sigma). Secondary antibodies conjugated with Alexa Fluor488, and Cy3 (Jackson ImmunoResearch) were used at 1:5000 dilution to visualize the primary antibodies. Coverslips were mounted using the VECTASHIELD fluorescence mounting media with or without DAPI (Vector Laboratories) and sealed using nail polish. Some tissue sections were counterstained with Hoechst dye instead of DAPI. Stained tissue sections were imaged on a Nikon A1 confocal microscope using NIS elements software for image acquisition.

### Electron Microscopy

Kidney tissues were perfusion fixed using 4% PFA and 2.5% glutaraldehyde in 0.1M Cacodylate solution. For scanning EM kidneys were sent for processing to Dr. Vincent Gattone at the Electron Microscopy Core of Indiana University School of Medicine. For TEM, kidney tissues were post-fixed with 0.5% osmium tetroxide + 0.5% potassium ferrocyanide for 1 h and incubated in 50 mM uranyl acetate in acetate buffer for 30 min. Dehydration of tissues was carried out using a series of ethanol solutions (50%, 70%, 90% and 100% for 15 min each). The tissues were infiltrated with a mixture of ethanol and araldite (2:1, 1:1, 1:2 ratios for 2 h each and 100% araldite overnight) and cured with a fresh araldite resin at 60°C for 48 h. Sections of 70 nm thickness were cut with an ultramicrotome (RMC Powertome XL). MDCK cells (~5×10^4^) were cultured on 12 mm transwell permeable supports (Corning costar 3460) for 5 days. The monolayer of cells were fixed with 2.5% glutaraldehyde in 0.1 M cacodylate solution for 20 min, and post-fixed with 1% osmium tetroxide for 30 min. Dehydration and embedding in araldite was carried out by following the same steps as above. Thin sections of 60 nm were cut parallel to the transwell membrane and the grid-mounted sections were stained with uranyl acetate followed by lead citrate. Images of cross-sections through cilia were taken on a JEOL JEM-2100 LaB_6_ transmission electron microscope.

### Cell culture

MDCK cells (ATCC) were grown in Eagle’s minimum essential medium with 10% fetal bovine serum and penicillin/streptomycin. Cultured cells were maintained in 95% O_2_/ 5% CO_2_ environment. For establishing stable lines of MDCK cells expressing dnMaml fused to EGFP^42^ or Akap12 fused to EGFP a kind gift from Irwin Gelman (Addgene plasmid #61294), cells were seeded in 24-well plates to reach 95% confluency the following day. Cells were transfected with neomycin selection expression vectors with Lipofectamine 2000 (Invitrogen) and GFP positive clones were isolated under selection and expanded. NIH3T3 cells were plated on fluoro dish (WPI) and transfected with 5HT6 fused with GFP, a kind gift from Kirk Mykytyn (Addgene plasmid # 35624)^43^ to visualize cilia. The cells were serum starved for 36 hours to promote exit from the cell cycle and initiate ciliogenesis. During the 36 hours cells were treated with DMSO or 10μm DAPT, and DAPT was added every 12 hours prior to live cell imaging using Nikon A1 confocal microscope.

#### Reporter assay

Cells were plated in 24-well plates and allowed to reach 90% confluency prior to transfection with empty expression vector or NICD expression vector along with TP-1 reporter and a β-galactosidase expression vector. Cells were lysed 48h post transfection and firefly-luciferase activity and β-galactosidase activity were determined. Assays were performed in triplicate and firefly luciferase activity was normalized to β-galactosidase activity.

#### 3D spheroid/single lumen formation assay

Low passage MDCK cells were used for lumen formation assay in 3D. One day prior to starting lumen formation, a confluent dish of MDCK cells was passaged at 1:10 to have the cells at their highest growth and proliferative stage. On the day of the experiment, freshly trypsinized cells were resuspended at a density of 3×10^4^ cells/ml in collagen solution (24mM Glutamine, 2.35mg/ml NaHCO_3_, 1X MEM, 20mM HEPES pH7.6, 2mg/ml collagen I in a total volume of 6.255ml). Cells were then plated in chamber slides and the collagen allowed to solidify in a 37C oven in absence of CO_2_. Cells were allowed to grow in this collagen matrix for 5 days. For immunostaining, the collagen matrix was digested in collagenase-1 (Type CVII, 1000U/ml) in PBS+ (PBS with 0.9mM Ca^2+^ and 0.5mM Mg^2+^) for 10min at RT. Cells were fixed in 4%PFA for 30min at RT, washed 3 times with PBS+ for 10min each. Cells were made permeable by incubation with PBS containing 10%FBS and 0.5% TritonX-100 for 30min at RT. Phalloidin (at 1:10,000 dilution, P1951, Sigma) and anti-β-catenin (at 1:500 BD Transduction Labs) were incubated with the cells overnight at 4C. Cells were then washed 3 times in PBS+ prior to incubation with secondary antibody (1:400 dilution) and Hoechst (1:1000) for 2hr at RT. Cells were finally given 3 quick washes followed by 3 washes for 15min each with PBS+ and mounted using vectashield mounting medium without DAPI.

#### Polarized, quiescent MDCK cells and mouse kidneys were analyzed for primary cilia length

For ciliogenesis, MDCK cells were plated on 10μm thick polycarbonate transwell membranes inserts (Corning) with growth media in both upper and lower chambers. The day after plating, media was removed from the upper chamber to allow cells to grow in an air-liquid interface for up to 10 days. At the end of incubation, membranes were washed twice with PBS, fixed in 4% PFA for 10min followed by a 10min fixation in ice-cold methanol at −20°C. Membranes were then washed in PBS, blocked for 1hr at RT in blocking buffer containing 1% bovine serum albumin (BSA), 0.2% powdered skim milk, and 0.3% Triton X-100 in PBS, and then incubated with acetylated alpha tubulin (Sigma, T7451, 1:500 dilution in blocking buffer) overnight at 4°C. Following extensive washes with PBS, the membranes were incubated with Alexa488 conjugated secondary antibody (1:5000) along with gamma-tubulin (Sigma, T7604, 1:200) for 1hr at RT. Membranes were mounted in Vectashield mounting medium containing DAPI. Confocal images of ciliated cells were captured using Nikon A1 confocal microscope and cilia length from each cell was measured using the Annotations and Measurements tool of the NIS- Elements Software (Nikon). Similarly, three week old mouse kidneys from *Rarb2-Cre;RBPJ ^f/f^* and control littermates were sectioned, stained for megalin and acetylated alpha tubulin, imaged using Nikon Ni-E microscope and cilia length from each cell was measured using the Annotations and Measurements tool of the NIS- Elements Software (Nikon).

### Quantitative PCR

Flash frozen whole mouse kidneys were homogenized using a tissue homogenizer (Biospec Products, Inc.). Total RNA was extracted from the whole kidney lysate using RNeasy Midi-Kit (Qiagen) according to manufacturer’s instructions. One microgram of RNA was reverse transcribed using oligo dT with Goscript reverse transcription kit (Promega). For each primer pair standard curves were generated with serially diluted cDNA reverse transcribed from whole mouse kidneys to determine the efficiency of each primer pair. Each sample was measured in duplicate and the relative gene expression levels were normalized to that of *GAPDH*. Primer sequences are available upon request. The relative expression levels were determined using the ΔΔCt method.^44^

### Statistical analysis

Excel was used to perform two-tailed unpaired t-tests, after testing for equal variance between groups using the F-test unless stated otherwise. The resulting p values are stated in the text and/or figure legends. Scatter plots were generated using GraphPad Prism 5.

## Acknowledgements

Research reported here was supported by a grant from NIDDK of NIH under award number R01DK106135. We thank the Molecular Pathology Core, Imaging Core and Functional Genomics and Bioinformatics cores at Sanford Research for technical assistance. These Cores at Sanford Research were supported by NIH grants P20GM10358, P20GM121341 and P20GM103620.

## Supplementary Materials

**Figure S1.** Inhibition of Notch signaling in NIH3T3 cells increases primary cilia length. NIH3T3 cells were transiently transfected with 5HT6 (serotonin receptor) fused with GFP to visualize cilia. A &B. Serum starved DMSO treated NIH3T3 cell has a cilium (arrow in A) which is smaller in length compared with cilium on a serum starved DAPT treated NIH3T3 cell (arrow in B). C. Average cilia length in DMSO treated cells is 6.03 +/- 2.48 μm, which is significantly less than the average cilia length of 17.36 +/- 10.85 μm in DAPT treated cells. * denotes p<0.05, student t-test n=23 cilia per treatment. Scale bars are 5μm.

**Figure S2.** Notch signaling deficient kidney epithelial cells have long primary cilia. Kidneys from three week old mice were stained with megalin, a proximal tubule marker, and acetylated-alpha-tubulin to visualize cilia. A. Wild type kidneys have a single apically located primary cilia (arrow) which are on average 4.2 +/- 1.1 μm. B. Notch-signaling-deficient kidney epithelial cells have long primary cilia (arrows in B & D) with an average length of 8.4 +/- 0.6μm. C. The cilia length is significantly longer in Notch-signaling-deficient kidney epithelial cells (MT) compared with wild type kidney epithelial cells (WT). * denotes p<0.05, student t-test n>32 cilia per kidney, and three kidneys per genotype. Scale bars are 10μm.

**Figure S3.** Notch signaling deficient kidney epithelial cells occasionally have supernumerary centrioles. Wild type (WT) MDCK cells (A &B) have two mitotic spindle poles, while dnMaml expressing MDCK cells have abnormal increase in number of centrosome (C&D) sometimes resulting in multiple mitotic spindle poles and likely aberrant segregation of genomic DNA. Examination of centrioles in mouse kidneys revealed at most two pairs of centrioles per epithelial cells in WT kidneys (E), whereas some cyst-lining epithelial cells (arrow in F) of Notch-signaling-deficient kidneys.

