## Supplementary figures and images for "Notch signaling regulates *Akap12* expression and primary cilia length during renal tubule morphogenesis"

### Supplemental Figure 1

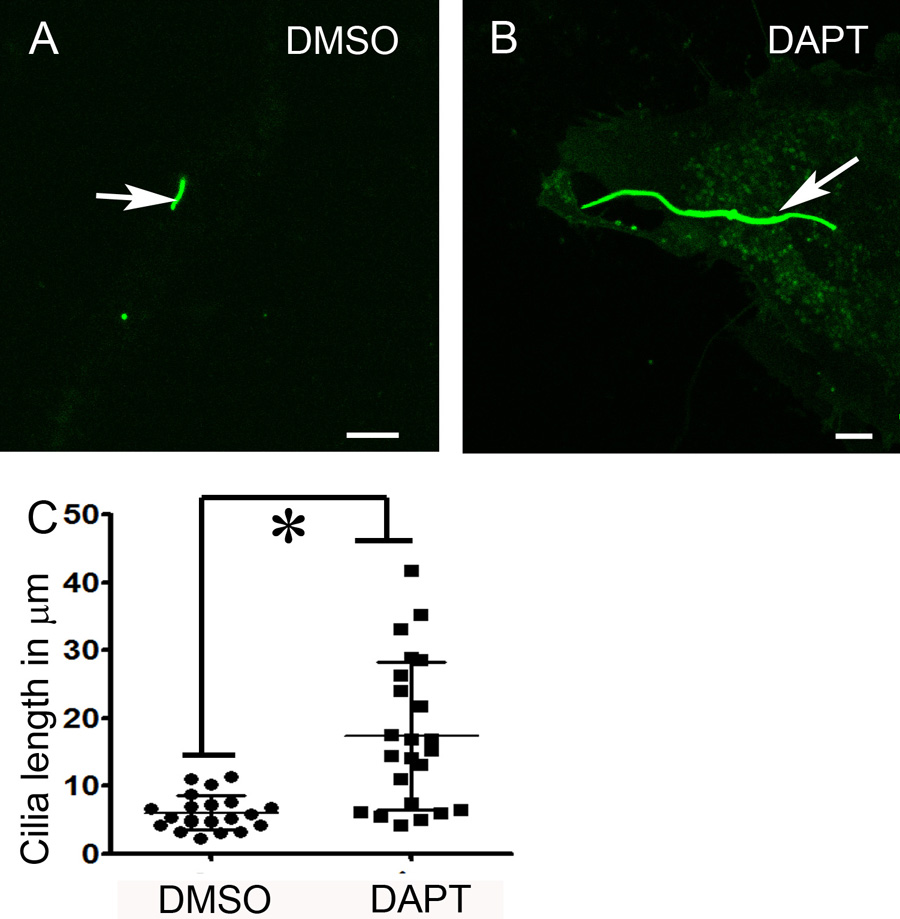

### Supplemental Figure 2

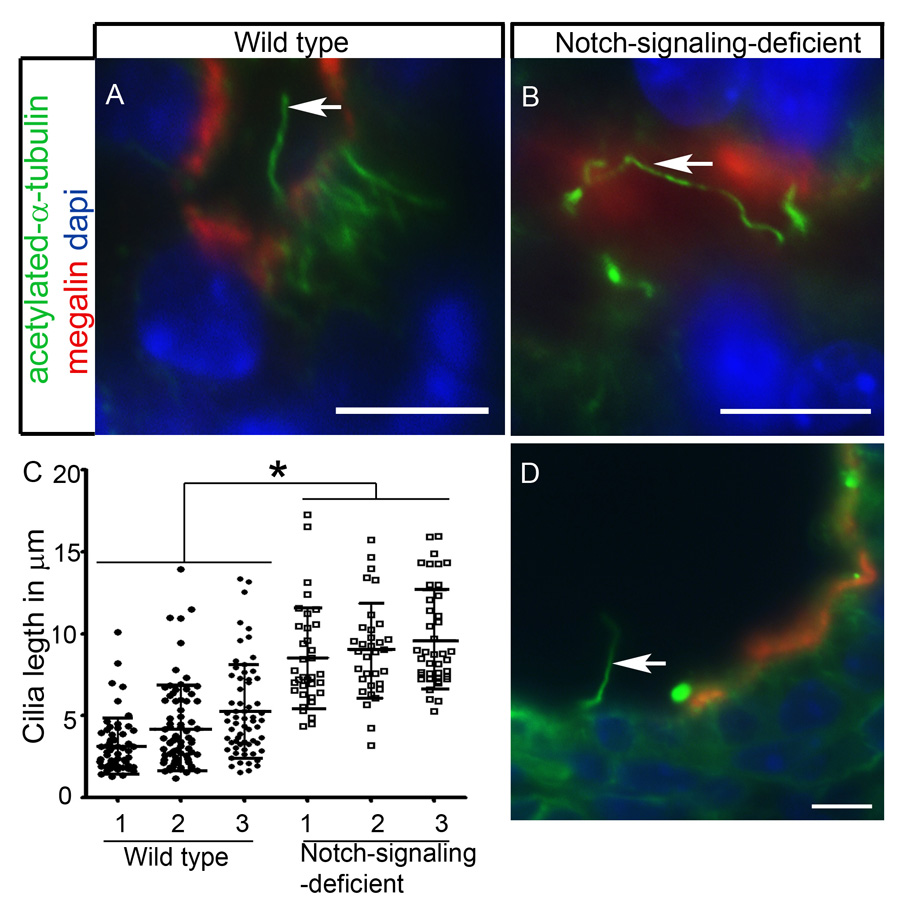

### Supplemental Figure 3

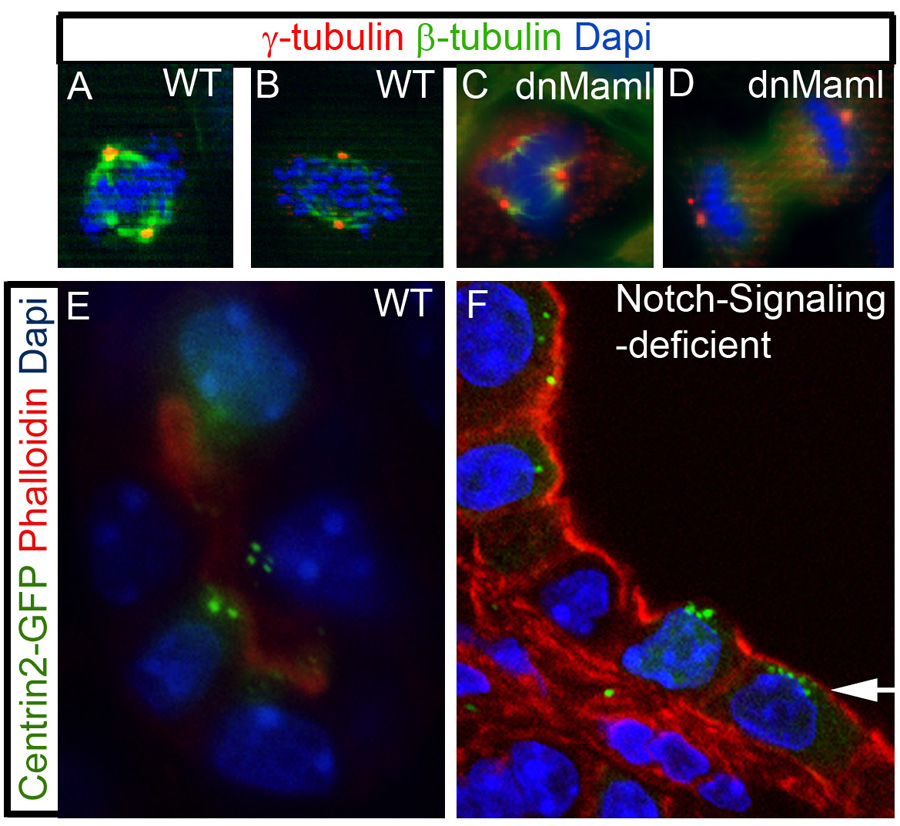
